# Health care service providers’ experiences, understanding and conceptions of voluntary medical male circumcision in KwaZulu-Natal, South Africa

**DOI:** 10.1101/753046

**Authors:** Celenkosini Thembelenkosini Nxumalo, Gugu Gladness Mchunu

**Affiliations:** KZN Department of Health, Ndwedwe Community Health Centre, South Africa; Discipline of Nursing, School of Nursing and Public Health, University of KwaZulu-Natal, South Africa

**Author notes:** Correspondence: 1. Celenkosini Thembelenkosini Nxumalo, KZN Department of Health, Ndwedwe Community Health Centre, Private Bag X528 Ndwedwe 4342, South Africa.

## Abstract

**Background:** There is compelling evidence that voluntary medical male circumcision reduces the chances of heterosexual transmission of HIV infection. Health care workers are among the key influencers in terms of scale up of VMMC as they are often involved in mobilization for uptake. Adequate knowledge and competence are essential to ensuring that the delivery of VMMC services is line with the recommended comprehensive package of HIV prevention services.

**Aim:** The aim of this study was to analyse health care service providers’ conceptions, experiences and understanding of VMMC in KwaZulu-Natal, South Africa.

**Methods:** The study employed a qualitative approach using a phenomenographic design. Ethical clearance to conduct the study was obtained from the University of KwaZulu-Natal’s Biomedical Research Ethics Committee (BE627/18).Data were collected from a purposive sample of 15 health care worker who were doctors, nurses and clinical associates working in six different rural clinics in KwaZulu-Natal, South Africa. Individual in-depth interviews were used collect data guided by a semi-structured interview schedule. An audiotape was used to record the interviews, which were then transcribed verbatim, and analysed thematically.

**Results:** Categories of description in healthcare providers’ experiences, conceptions and understanding of voluntary medical male circumcision emerged. The findings of this study revealed that health care workers conceptions, experiences and understanding of VMMC were influenced by stereotypical cultural, religious and traditional beliefs. The challenges of implementing VMMC were shortage of staff and poor training of health care workers on VMMC.

**Conclusion:** Tailored messaging targeting health care workers misconceptions and poor understanding of VMMC in necessary. In addition, resource allocation for training and infrastructure could significantly improve the quality of VMMC services and uptake thereof.

## INTRODUCTION

The results of three Randomized Controlled Trials conducted in Kenya, Uganda and South Africa have proven that Voluntary Medical Male Circumcision (VMMC) is an effective biological HIV prevention strategy (1–3). Following these results, the World Health Organisation (WHO) recommended that regions of high HIV prevalence adopt VMMC as an additional HIV prevention intervention (4). In response to these recommendations, VMMC was introduced in KwaZulu-Natal (KZN), South Africa.

Since the roll-out of VMMC, health care workers have been at the forefront of implementing the service as stipulated by standardized protocols and procedures (5, 6). Adequate clinical training, knowledge and competence are some of the essential elements to ensuring delivery of an efficient and effective VMMC service, particularly in the context of the current model of health service delivery in South Africa (7).

The implementation of VMMC in KZN, South Africa has been robust with 1.2 million medical circumcisions done since 2010. A significant proportion of these medical circumcisions have been on boys in the 10-14 year age group, allowing for long-term protection of this age group against HIV infection (8, 9). However, if the province and country is to yield immediate curbing of new infections, targeting males in the sexually active 18-49 age group becomes of paramount importance.

Research reviewed on the uptake of VMMC reveals that there are several challenges that hinder uptake among males. Among these are individual perceptual factors, social influencers such as peers, key figures and role models, and structural barriers pertaining to availability health resources to meet the needs of men (10–12). The scale up of VMMC so that stipulated targets are met is also dependent upon health service providers’ attitudes, perceptions and beliefs about the efficacy of VMMC and may influence how health workers mobilize for scaling up of VMMC.

While VMMC has been implemented in KZN, there is a paucity of data concerning it, particularly with reference to the perceptions and experiences of health care workers regarding the service. Data on the perceptions of VMMC currently appears to be limited to the perceptions of pharmacy and nursing students(13). Research on health care workers’ attitudes and understanding of VMMC emanates from a time prior to the roll-out of VMMC in South Africa (14), while current data on health care workers’ attitudes and perceptions of implementing VMMC are limited to two regions in Africa, both of which are urban settings(15, 16).

In addition, there is a dearth of data capable of providing an in-depth understanding of how VMMC is understood, experienced and conceived, particularly by health care providers within a context, which is heavily influenced by culture, tradition and notions of gender norms of patriarchal hegemonic masculinity, especially with regard to medical circumcision.

The purpose of the present study is therefore to analyse health care service providers’ experiences, understanding and conceptions of VMMC in KZN, South Africa. An awareness of health care workers’ experiences, understanding and conceptions of VMMC within a socio-cultural context has important implications for policy in terms of education and training regarding VMMC and quality improvement relating to how services are rendered; which may potentially influence the uptake.

## AIM

To analyse health care service providers’ experiences, understanding and conceptions of VMMC in KZN, South Africa.

## RESEARCH QUESTION

1. What are the health care service providers’ experiences, understanding and conceptions of VMMC in KZN, South Africa?

## RESEARCH DESIGN

A qualitative approach using phenomenographic study design was used. This study was part of a larger study conducted to analyse the conceptions, experiences and understanding of primary health care stakeholders’ regarding VMMC in order to develop a relevant intervention to support scale up of VMMC in KZN, South Africa.

Phenomenography, is a qualitative research design that is interpretive in nature and seeks to describe, discern and analyse the ways in which individuals conceive, understand and experience a phenomenon in the world around them. It does not consider the subject and aspect of the world as separate entities, but rather the individual’s experience, conception or understanding are seen as setting up a relation between that person and a given phenomenon in the world.

## RESEARCH SETTING

This study was conducted at six selected rural primary health care facilities in six different health districts offering VMMC services in KZN, South Africa.

## RESEARCH POPULATION

Health care workers, namely, all categories of nurses, doctors and clinical associates working at the selected primary health care facilities, formed part of the study. The clinicians interviewed included those who were directly involved in rendering VMMC services at the selected facilities.

## SAMPLING METHOD

Selection of health districts and facilities was done purposively. Theoretical sampling was used to select participants to form part of the study. Data were collected from a sample of 15 health care service providers.

## INCLUSION CRITERIA

- Health care workers at primary health care level namely nurses, doctors and clinical associates directly involved with VMMC.

## DATA COLLECTION

A semi-structured interview guide was used to collect data from the target population by means of an in-depth individual interview recorded using an audiotape.

Data collection took place after ethical approval from the University of KZN’s Biomedical Research Ethics committee (BE627/18) as well as the provincial department of health. Interviews were conducted after informed consent was obtained from the participants and lasted for approximately 45 minutes.

## DATA ANALYSIS

After data collection, transcription was done and data were analysed thematically. The steps of phenomenographic data analysis were followed. They are:

1. *Familiarisation step*: Transcripts were read to become familiar with their contents.
2. *Compilation step*: This step required focused reading in order to deduce differences and similarities from the transcripts. Through this process the researcher was able to identify the key elements in answers.
3. *Condensation step*: Extracts that seemed to be meaningful and relevant to the study were extrapolated.
4. *Preliminary grouping step*: The focus was on locating and classifying similar responses into preliminary groups.
5. *Preliminary comparison of categories*: This step entailed the revision of the initial list of categories to bring forth comparisons among the preliminary listed categories.
6. *Naming the categories*: Following confirmation of the categories, they were named to emphasize their essence.
7. *Final outcome space*: In the final step, the aim was to discover the final outcome space based on their internal relationships and the ways of understanding, conceiving and experiencing the phenomena.

## TRUSTWORTHINESS

### 1) Credibility

Audiotapes were used to collect data, which was then transcribed verbatim and then analysed thematically. The data analysis was done under close supervision by the researcher’s supervisor.

### 2) Dependability

The researcher transcribed the audio data and thereafter read the data to ensure that it matched what was on the audio tape.

### 3) Confirmability

The help of an experienced qualitative researcher was sought when the data were analysed.

### 4) Transferability

This was ensured by purposively selecting the data collection sites and theoretically sampling the participants facilitate transferability.

## RESULTS

Fifteen participants formed part of the study. Most of the participants were registered nurses who had been involved with VMMC for at least two years. Demographic details of the participants are detailed in table1.

### Conceptions of VMMC

#### a) Foreskin removal

Health care workers in this study seemed to be convinced that VMMC is the removal of the foreskin of the penis of a male, the extent of foreskin removal is, however not explicitly mentioned. This removal is, however seen to be medically beneficial. This is justified by the following statements:

> *“A surgical procedure where the foreskin of the male genitalia is removed as it is a good medium of transmission for HIV and STI’s.”* (Participant 3)
>
> *“Medical male circumcision is the removal of the foreskin, a small piece of the skin. They do not remove the whole thing. This is to help prevent contracting any diseases. You are less susceptible to diseases if you are circumcised.”* (Participant 1)
>
> *“In short I would say it is removal of the foreskin covering the glans. It’s so that the glans is exposed.”* (Participant 2)
>
> *“Medical male circumcision is when the foreskin is removed from a boy’s penis.”* (Participant 7)

#### b) A procedure for ‘boys’

It was found that while health care workers may be aware that medical circumcision is for men of all age groups, it appears that a significant number of them believe that this procedure is mainly for young boys. This was noted in the practice of implementing VMMC and their narration of their opinions about VMMC and the best time to perform medical circumcision. The excerpts below serve to support this category:

> *“For convenience, I think it is better circumcise boys starting from eight years and above…Here we do them when they are that age because they are able to tolerate anesthesia.”* (Participant 4)
>
> *“The older generation of boys and men are a problem because they often have adhesions that are a challenge when doing circumcision. In boys of a very small age, there are also challenges so I think 12 years is okay.”* (Participant 1)
>
> *“It is better to do the procedure on boys from 12 years old because they understand and know everything better. When you explain things to them, they do not have a problem only when you inject them there is some bit of challenge but after that they are fine and you carry on.”* (Participant 5)
>
> *“People here are not interested in being circumcised, especially the older ones… most of the time we have children.”* (Participant 8)

#### c) Service to be rendered by men

Most health care workers believe that since VMMC has deep cultural and religious connotations underpinned by notions of traditional hegemonic masculinity bias, health care workers rendering such services should preferably be men. This opinion stemmed from their experience rendering VMMC services and interacting with men at all levels of care. The following statements support this category:

> *In my opinion, I can say that sometimes you do not become comfortable to be assisted by a female. (Participant 7)*
>
> *From my observation most men are not happy when it is women who are involved in doing the MMC, sometimes some of them just turn away in the last moment when they see women doing it, but school children normally don’t have a problem. (Participant 1)*
>
> *In the rendering services, I think it best for a man to be performing the circumcision seeing that the procedure is very much influenced [by] traditional beliefs especially here in the Zulu culture. Even more this is something private, for a women to be involved seems sort of disrespectful to a man and may cause him to not want to get medically circumcised. (Participant 4)*
>
> *In terms of performing the procedure, it must be males. (Participant 14)*

### Experiences of VMMC in KZN

#### a) Walk-in VMMC services and MMC camps

Health care service providers reported two approaches of implementing VMMC: walk-in VMMC services and provision through camps. The walk-in services are rendered by personnel employed by the department of health and occurred daily. Walk-in services are provided when a male comes into the facility and voluntarily requests MMC services. The camp system normally occurs once a month and involves intense mobilization by personnel working for the department of health and a private Organisation working with the department of health.

> *“We have community caregivers that market it in the community, both for daily MMCs and the camps we host once a month.” (Participant 7)*
>
> *“We are usually mobilizing and using the camp system… we are mainly getting young ones at primary school level, those that are in high school come too but not as much.” (Participant 2)*
>
> *“We do same day (walk-INS) but it depends on that person’s status. There is a procedure to follow when one tests positive. If they are positive, we also follow the appropriate procedure.” (Participant 3)*
>
> *“We camp monthly, then you find kids are at schools so sometimes we go there to mobilize.” (Participant 9)*

#### b) Poor training and preparation with service delivery shortfalls

The lack of sufficient training and guidance with regard to the clinical practice of medical circumcision was widely reported by participants who also stated that they had received no clear guidance as to what their role was and what was expected of them. In addition to poor practical training, participants reported problems with the organization of the program at an operational level and also on the side of policy and strategic management. Participants reported that the generally poor way in which the programme had been managed could also account for the programme’s poor uptake in the immediate target age group. The following accounts of participants’ experiences of VMMC give an account of this:

> *“The doctor here leaves at 11 AM. We remain here but are unable to do anything… sometimes he does not come to the clinic at all because of different things. All bookings done on the day he decides not to come get cancelled and rebooked again and sometimes you lose the initial number of males you had.”* (Participant 2)
>
> *“A huge challenge is that we are not trained…”* (Participant 6)
>
> *“If you book them, they don’t come. Some even when you rebook and tell them to come early they still end up not turning up the next time*. (Participant 9)
>
> *“Our district has mostly deep rural areas and people cannot come to us to circumcise because of distance and the transport challenge that comes with it.” (Participant 12)*
>
> *“We do not have all personnel…. we do not have a nursing assistant and enrolled nurse, we do whatever we can but it is difficult.”* (Participant 5)

#### c) Misconceptions regarding VMMC

Most participants reported that despite concerted efforts to educate men about medical circumcision and the benefits thereof, adverse side effects related to postoperative infections and misconceptions and negative stereotypical beliefs often spread easily among community members, especially when someone who had undergone the procedure at a health institution relayed such information. The participants seem to concur with one another in that negative information is seemingly easily absorbed and acted upon, contributing to the low uptake of VMMC. The following statements supported this:

> *“They have a belief that after MMC you can have a reduced private part.”* (Participant 14)
>
> *“In the older age group it seems that the problem is if you perform this thing you have to abstain from doing marital obligations for two weeks or more and this is not a pleasant experience. I also personally have not been circumcised because when I imagine myself refraining now from doing this thing for two weeks, I am going to lose my mind. And in that two weeks’ time there is no way that I cannot think of… [Sex] as my mind thinks of that the stitches will burst. I would get an erection and it will be quite an event you see. Those are the things that cross my mind every time.”* (Participant 4)
>
> *“…something can go wrong during the procedure and that wrong may steal your joy forever, so why expose yourself to the risk when you had no problem?”* (Participant 10)

### Understanding of VMMC

#### a) HIV and STI prevention

Most participants seemed to understand that VMMC reduces the chances of HIV infection. They were also aware that medical circumcision only partially reduced the chances of infection. However, some participants do not understand the degree of such reduction.

> *“When you are uncircumcised, there are infections on the collar of the penis.”* (Participant 9)
>
> *“Medical circumcision is helpful to both males and females in that once the foreskin is removed there is reduced chances of infections like HPV, which causes cancer in females, and with males, also there is reduced incidence of other STIs and even penile cancer.”* (Participant 13)
>
> *“Circumcision helps to reduce the chances of contracting HIV and other sexually transmitted diseases by 60 per cent.”* (Participant 11)

#### b) Traditional verses medical circumcision

A significant number of participants seem to think about traditional circumcision in terms of the complications associated with the procedure due to the manner and context in which it is carried out. Participants thus seem to understand that medical circumcision is a safer option compared to the traditional method. This could be attributed to the reported adverse events that have occurred in traditionally circumcising communities. The understanding of health care workers regarding medical and traditional approaches to circumcision seems to be largely influenced by their clinical orientation to health care, including medical circumcision.

> *“I know more about medical circumcision because that is what I have been doing here. It’s not painful, they inject you and give you medication. As far as I know, at the mountain they don’t give you any treatment. It is not safe because you can end up with infections. You can end up dead if you do not receive any treatment.”* (Participant 15)
>
> *“I don’t know about traditional circumcision but I know we have had so many problems. People dying, especially in the Eastern Cape, because of issues of hygiene and sterility.”* (Participant 11)
>
> *“Overall, medical circumcision is safe….I’m not sure if they even deal with the inner prepuce at the mountain.”* (Participant 3)

#### c) Healing time versus circumcision time

In terms of healing time, the consensus among participants is that it takes six weeks for a male to be fully healed after VMMC. However, in terms of the time at which medical circumcision can be performed, participants appear to be divided. Some believe that as early as birth would be the right time, while others reported anywhere from 8 to 12 years. The reasons for such understanding stem from the perception that if males are circumcised while too young, they will not be able to withstand anesthesia while others feel that young boys will not generally cope well with the procedure.

> *“It takes 6 weeks, we tell them that. We also teach them that after two weeks, maybe they will think that the wound is healing. But that does not necessarily mean that it is completely healed to engage in sexual intercourse… they need to wait for 6 weeks.”* (Participant 1)
>
> *“There is no specific time, but for convenience, it’s better to do them when born. But because of the anesthesia challenges it’s better to do them when they are 8 and above because they are better able to tolerate the procedure…”* (Participant 4)
>
> *“For us, we start at 12 because most of the time, the younger ones less than 10 cry a lot. You cannot work properly. You cannot do the procedure properly because they will give you a little trouble. They cry and cannot keep their hands away from the procedure area. It takes even longer to finish doing them.”* (Participant 3)
>
> *“In my understanding there is no stipulated time suitable for undergoing circumcision, a male child can get circumcised immediately after birth if their parents’ consent to it.”* (Participant 13)

## DISCUSSION

The present study analysed health care services providers’ experiences, understanding and conceptions of VMMC in KZN.

Health care service providers’ experiences of VMMC centered on the implementation and delivery of VMMC services within the primary health care context. Participants reported that VMMC was rendered through two approaches: 1) Walk-in VMMC services, 2) VMMC camps. In the Iringa region of Tanzania, it was found that the camp approach was used to provide quality high volume VMMC services (17)

VMMC would typically entail basic health screening, health education and obtaining consent. The current model of health service delivery particularly in South Africa calls for the adoption of promotive and preventive strategies (18) meaning that VMMC should typically be rendered within an array of comprehensive HIV prevention services.

Challenges with implementing VMMC particularly at primary care level seem to be mainly service delivery related and this stemmed from a shortage of both human resources and, in certain cases, insufficient infrastructure to perform the service. The shortage of staff is a global phenomenon cited in many international studies (19–21).

The shortage of staff is a major barrier to productivity and compromises quality of health care (22–24). These challenges were also cited in Kenya and were addressed through specific changes in policy and the structure of health facilities(6).

Lack of training of staff working permanently at the clinics, and generally poor acceptance of VMMC by males, particularly those in the older age groups were also challenges to implementation. Similarly in a study to explore beliefs and attitudes of health workers regarding medical circumcision in Haiti, it was found that while levels of knowledge and positive attitudes concerning VMMC were high, lack of adequate training among physicians and doctors was a challenge to service implementation (15). Multi-sectoral collaboration in such instances has been found to be useful as it leverages more knowledge and expertise as a result of combined strengths working towards achieving a common goal (25).

Participants’ understanding of VMMC was related to the partial protection of VMMC against HIV infection and certain sexually transmitted infections. Health care workers were also aware of the indirect benefits of VMMC to female clients. In kenya women also demonstrated good understanding of the partial protection of VMMC (26). In an earlier study conducted in South Africa similar finding were also found among women (27)

While many health providers were aware of the nature of VMMC, most were unclear about the manner in which traditional male circumcision was undertaken. However, most perceived medical circumcision to be safer than traditional circumcision. Similar findings were cited in Malawi among circumcising and non-circumcising communities (28)

Participants seemed to understand the VMMC healing process and six week sexual abstinence period, however, in terms of performance of VMMC, most seemed to think that younger boys would benefit more from VMMC because of the perception that younger boys would be less likely to have postoperative complications. This indicates a misperception on VMMC on the part of health care workers. These findings concur with the belief that intensification of neonatal circumcision is better because it is associated with fewer medical risks (29). Other studies have mainly sited poor knowledge of health care workers in relation to VMMC and the need for HIV testing (30).

In terms of delivery of the service, health care service providers were of the opinion that VMMC should be rendered by males only. This perception highlights the influence of gender norms of masculinity on VMMC. In KwaZulu-Natal, South Africa it was found that VMMC is associated with perceptions of gender identity and masculinity including sexual performance, sexual image and self-identity (31). In Zimbabwe the perceived challenge to masculinity was one of the major barriers to acceptability and uptake of VMMC (32).

## LIMITATIONS

The study was limited in that only fifteen participants from only six different health district formed part of the study. Therefore, the results may not necessarily be generalizable to all settings. Another factor is the methodology used which does not allow data to be quantified. Nevertheless, the results provide a deeper awareness of the experiences, conceptions and understanding of VMMC within a rural health context.

## CONCLUSION

The study findings suggest that the experiences, understanding and conceptions of health care service providers are influenced by an array of factors. The experiences of these health care workers were related to implementation challenges stemming from lack of training and insufficient infrastructure, together with a poor acceptance of VMMC by males. Participants’ understanding of the procedure was influenced by their culture and tradition together with general misinformation about the procedure and the biology of the penis and foreskin. The conceptions of health care workers also seemed to be influenced mainly by stereotypical norms of culture, tradition and the prevailing hegemonic masculinity.

The study findings provide important information on some of the impediments to successful scale up of VMMC. Moreover, they highlight the need for training interventions to be directed towards health care workers in order to address misinformation about VMMC, such interventions should take cognizance of the traditional and religious issues related to medical circumcision.

Greater support is needed in the form of resource allocation for VMMC to ensure provision of adequate and skilled health personnel to provide quality VMMC services.

## Acknowledgements

The research reported in this publication was supported by the Fogarty International Center (FIC), NIH Common Fund, Office of Strategic Coordination, Office of the Director (OD/ OSC/CF/NIH), Office of AIDS Research, Office of the Director (OAR/NIH), National Institute of Mental Health (NIMH/NIH) of the National Institutes of Health under Award Number D43TW010131. The content is solely the responsibility of the authors and does not necessarily represent the official views of the National Institutes of Health.

